# Rapid and scalable preparation of bacterial lysates for cell-free gene expression

**DOI:** 10.1101/162768

**Authors:** Andriy Didovyk, Taishi Tonooka, Lev Tsimring, Jeff Hasty

**Affiliations:** BioCircuits Institute, University of California San Diego, La Jolla, CA 92093, USA; San Diego Center for Systems Biology, University of California San Diego, La Jolla, CA 92093, USA; Department of Bioengineering, University of California San Diego, La Jolla CA, USA; Molecular Biology Section, Division of Biological Science, University of California San Diego, La Jolla, CA 92093, USA

**Keywords:** cell-free gene expression, *in vitro* transcription-translation, cell lysates, autolysis, biosensors

## Abstract

Cell-free gene expression systems are emerging as an important platform for a diverse range of synthetic biology and biotechnology applications, including production of robust field-ready biosensors. Here, we combine programmed cellular autolysis with a freeze-thaw or freeze-dry cycle to create a practical, reproducible, and a labor- and cost-effective approach for rapid production of bacterial lysates for cell-free gene expression. Using this method, robust and highly active bacterial cell lysates can be produced without specialized equipment at a wide range of scales, making cell-free gene expression easily and broadly accessible. Moreover, live autolysis strain can be freeze-dried directly and subsequently lysed upon rehydration to produce active lysate. We demonstrate the utility of autolysates for synthetic biology by regulating protein production and degradation, implementing quorum sensing, and showing quantitative protection of linear DNA templates by GamS protein. To allow versatile and sensitive β-galactosidase (LacZ) based readout we produce autolysates with no detectable background LacZ activity and use them to produce sensitive mercury(II) biosensors with LacZ-mediated colorimetric and fluorescent outputs. The autolysis approach can facilitate wider adoption of cell-free technology for cell-free gene expression as well as other synthetic biology and biotechnology applications, such as metabolic engineering, natural product biosynthesis, or proteomics.

## 1 Introduction

Cell-free gene expression (CFE) has seen a tremendous surge in usage in biomedical research and biotechnology (*1*–*4*). Novel CFE applications include building field-ready biosensors (*5*), including Zika and Ebola virus sensors (*6*), producing biomolecules including natural products for therapeutic and research purposes (*7*–*10*), prototyping synthetic gene circuits (*11*, *12*), developing synthetic cells (*13*). The basis of most CFE systems is lysate derived from living cells. *E. coli* cell lysates are extensively used because of their high efficiency of expression from bacterial and phage promoters and because of the availability of a large set of well-characterized molecular components and genetic tools. However, production of bacterial cell lysates, especially retaining high transcriptional activity from endogenous promoters, presents a major bottleneck to wider adoption of CFE systems in synthetic biology and biotechnology. Traditionally, the established preparation procedures require complex protocols, special experience, and expensive equipment for mechanical cell disruption (French press or bead beater) that is not readily available in most laboratories and that limits the throughput in terms of sample size and numbers (reviewed in ref. (*14*–*16*)). Achieving reproducible results using these methods also requires considerable experience, limiting access to the technology. Recently, ultrasonication-based methods of lysate preparation have been developed to significantly reduce these constraints (*14*, *16*, *17*). Although these methods use less expensive equipment for cell disruption, they require careful thermal and power control to avoid protein denaturation during the sonication and are limited in sample size and numbers by the sonication hardware (*15*). To further address these limitations, a method of producing *E. coli* cell lysates that employs biochemical, instead of mechanical, cell disruption has been recently proposed (*15*). This method relies on degrading *E. coli* cell wall by externally added egg white lysozyme followed by cell disruption with hypotonic treatment. This method (further referred to as LoFT) eliminates the need for mechanical disruption equipment, addressing the accessibility and scalability issues. However, the method has a few potential limitations. First, it requires careful manual handling of mechanically sensitive lysozyme treated cells during the procedure to avoid premature cell lysis, that could lead to variability between batches or operators, and could limit sample throughput. Second, linear DNA, that is often used to accelerate and scale up template preparation for cell-free expression(*18*, *19*), is poorly protected in LoFT lysates, with close to an order of magnitude reduction of expression yields from *E. coli* σ70 promoters from linear versus circular DNA (*15*). Finally, to achieve the optimal LoFT lysate activity, a buffer exchange by dilution/concentration or dialysis is required (*15*), limiting throughout as well as leading to potentially detrimental loss of low molecular weight components from the solution (for example the precursors needed for N-acyl-homoserine lactone (AHL) synthesis in quorum sensing).

To address the aforementioned limitations, we present a robust and accessible method for *E. coli* cell lysate preparation with efficient expression from native *E. coli* promoters that requires only a single essential step – a freeze-thaw cycle of the strain programmed for autolysis using phage lambda endolysin (alternatively, a freeze-dry and rehydration cycle as described below). In short, after fermentation and harvesting, the cells are subjected to a freeze-thaw cycle and the lysate is separated from insoluble material by centrifugation at 30,000 g. Only a regular −80°C freezer, a set of centrifuges, and a vortex mixer are required for the preparation; the single reagent needed is a common S30A buffer (50 mM Tris-HCl, 60 mM potassium glutamate, 14 mM magnesium glutamate, 2 mM dithiothreitol, pH 7.7).

To demonstrate the utility of autolysates in synthetic biology and biotechnology, we characterized the autolysate-based CFE systems in a number of common scenarios. First, linear template DNA in autolysates was quantitatively protected by GamS protein, without noticeably affecting expression efficiency. Second, autolysate-based CFE systems could be endowed with ClpXP-mediated specific protein degradation activity by overproducing a variant of ClpX unfoldase within the autolysis strain during fermentation. Third, autolysate-based CFE systems supported a commonly used quorum sensing system from *Vibrio fischeri*: N-acyl-homoserine lactone (AHL) could be both synthesized and sensed efficiently. Finally, we observed that autolysate-based CFEs support efficient expression of *E. coli* β-galactosidase (LacZ). Therefore, to enable sensitive, versatile, and inexpensive reaction readout using LacZ, we engineered *lacZ*-deficient autolysis strain that produces autolysates with no background LacZ activity. Using LacZ deficient autolysates, mercury (II) biosensors sensitive to at least 12.5 nM of Hg^2+^ with both colorimetric or fluorescent outputs have been implemented.

Cell-free expression systems, freeze-dried on paper or other supports, have been recently used to create robust biosensors and *in situ* bioproduction reagents capable of room temperature storage for extended periods of time (*5*–*7*, *20*). Upon rehydration, such freeze-dried CFE systems support transcription-translation reactions enabling biosensing or bioproduction programmed by the supplied synthetic gene circuits. In this paper we demonstrate that this can also be achieved by directly freeze-drying live bacterial cells programmed for autolysis together with required cell-free reaction constituents (aminoacids, nucleotides, energy regeneration system, etc). Autolysis strains, freeze-dried this way, can subsequently be lysed by simple rehydration, initiating cell-free transcription-translation reactions (Figure 1). Although not directly investigated in this paper, long-term room temperature storage of the freeze-dried autolysis strain and cell-free reaction constituents is feasible, based on the previous work on lyophilization of cell-free expression systems (*5*, *20*, *21*). This practical approach could lead to more straightforward and inexpensive production of biosensors and bioproduction reagents. To demonstrate the utility of this approach we have implemented a sensitive proof of concept mercury (II) sensor sensitive down to at least 25 nM Hg^2+^.

**Figure 1:**
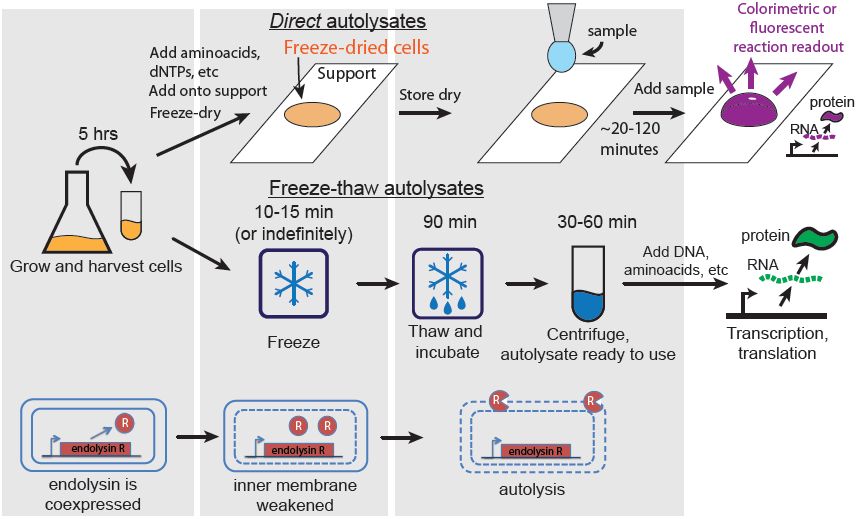
Overview of autolysate preparation. The autolysis plasmid constitutively produces low levels of phage lambda endolysin (gene R). The presence of endolysin has no significant effect on cell growth as long as the inner cell membrane is intact. These cells are then used to prepare cell autolysates either by freeze-drying and rehydration or by freeze-thawing (see the top and middle panels respectively). **(top)** Preparation of autolysates by freeze-drying followed by rehydration. Harvested cells are resuspended in a standard S30A *E. coli* transcription-translation (TX-TL) buffer with added standard TX-TL reaction constituents (amino acids, ribonucleotides, ATP regeneration system, etc). Optionally, gene circuit DNA can also be added in this step. The mixture is then freeze-dried on an appropriate support (*e.g.* paper, microtube, microwell plate). Upon rehydration, the cells lyse releasing the TX-TL machinery: the freeze-drying cycle compromises the inner membrane, and rehydration releases the endolysin protein to the periplasmic space which in turn lyses the cell. Gene circuit DNA and/or an analyte of interest can be added during rehydration. The reaction readout can be performed using fluorescent, colorimetric, or other output. **(middle)** Preparation of autolysates by freeze-thawing. Harvested cells are frozen in a standard S30A *E. coli* TX-TL buffer. Upon thawing the cells lyse. The cell extract is ready to use after 90 minute “run-off” incubation at 37°C and centrifugation to remove insoluble material. Transcription and translation are initiated upon addition of the desired plasmid DNA and standard TX-TL reaction constituents.

## 2 Results

### 2.1 Construction of autolysis plasmids

The key idea of our method is to lyse *E. coli* cells enzymatically using intracellularly produced lambda phage endolysin. Applying a freeze-thaw cycle compromises the inner cell membrane enabling endolysin to escape and disrupt the bacterial cell wall. In wild-type phage lambda, three genes S, R, and Rz are responsible for lysing the host cell (reviewed in (*22*)). Gene products R and Rz degrade the cell wall, while gene product S punctures the inner membrane of Gram-negative *E. coli* cells. Such disruption of the inner membrane is a prerequisite for R and Rz action; with the inner membrane intact high amounts of gene products R and Rz can accumulate without significant impact on cell growth (*23*). Cells containing R and Rz gene products, however, lyse efficiently when the inner membrane is disrupted chemically or by freezing-thawing the cells (*24*). Additionally, we demonstrate in this paper that freeze-drying followed by rehydration is also sufficient to release intracellularly produced gene product R and lyse cells. Thus, to create *E. coli* cells that efficiently lyse upon freeze-thawing or freeze-drying/rehydration, we engineered a plasmid that constitutively expresses low levels of gene product R (Figure 1 and Figure S1 of Supporting Information). We excluded the gene Rz, since it is not strictly essential for cell lysis (*25*).

### 2.2 Direct autolysate production by freeze-drying and rehydration *in situ*

We transformed the *E. coli* BL21-Gold (DE3) strain with the R gene containing plasmid. This strain lacks both major non-specific proteases (*lon, ompT*) and DNA endonuclease I (*endA*), thereby stabilizing proteins and plasmid DNA. The cells were grown in 2xYTPG medium (*26*), harvested by centrifugation, washed once with the standard lysate preparation buffer (S30A buffer) (*27*), and resuspended in the same buffer supplemented with reducing agent (Figure 1). To prepare autolysates by freeze-drying and rehydration, the harvested cells were mixed with the solution containing transcription-translation reaction constituents (amino acids, ribonucleotides, ATP, etc) and freeze-dried using a food-grade freeze-drier (Methods). After freeze-drying the mixture was stored dry and rehydrated with an appropriate amount of water supplemented with expression template DNA, to induce cell lysis and the transcription-translation reactions. To estimate the performance of the CFE systems produced in this way compared to the reference CFE system we measured the amount of green fluorescent protein (GFP) expressed from a strong lambda phage P_R_ promoter (*28*) (Figure 2A). GFP levels rapidly reached a plateau of approximately 3.4 μM within an hour with the maximum production rate reaching about 4.2 μM/hr (see the definition of the maximum production rate on Figure 1D, inset), using 15.4 nM GFP plasmid DNA. This is a relatively low activity compared to traditional *E. coli* lysates (compare to Figure 2B). Nevertheless, this low level of activity may already be sufficient for many applications, as demonstrated below for a mercury (II) biosensor. The described freeze-drying/rehydration method is very simple, but it leaves cell debris and genomic DNA in the lysate and this may be undesirable, for example leading to background expression of endogenous *E. coli* genes that could compete with the activity of the synthetic circuits of interest.

**Figure 2:**
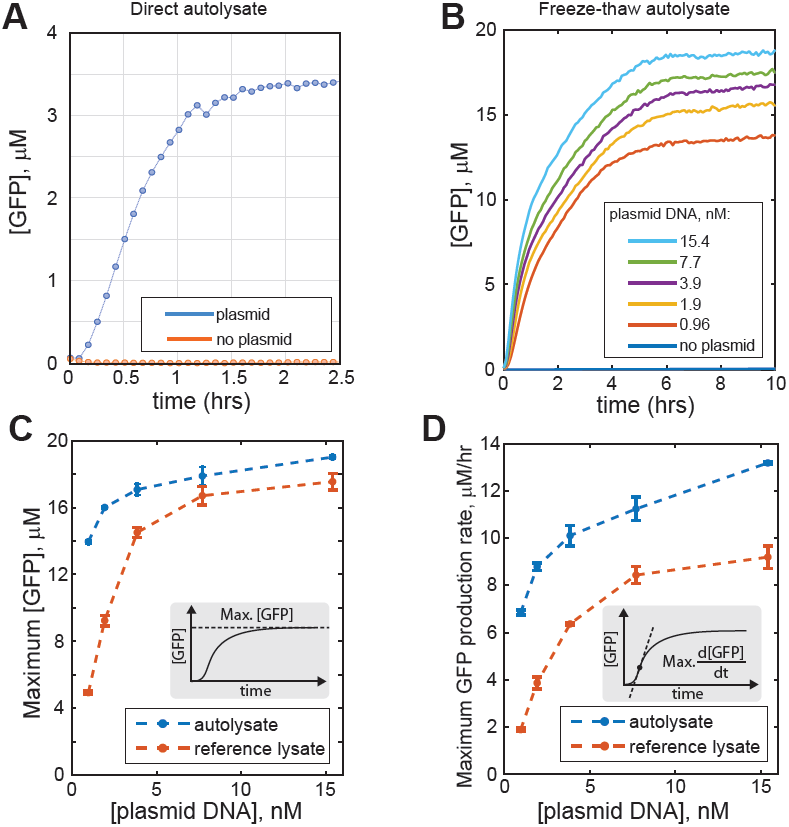
Characterization of gene expression in *E. coli* autolysates. (A) Time course of green fluorescent protein (GFP) synthesis in autolysate produced by rehydrating freeze-dried cells. (B) Time course of GFP production in freeze-thaw autolysate. Indicated concentrations of GFP producing plasmid DNA were added. (C, D) Autolysates produce comparable or higher levels of GFP and exhibit higher GFP production rates compared to the reference *E. coli* cell lysate.

### 2.3 Autolysate production by freeze-thawing

To produce traditional cell debris-free *E. coli* cell lysates using a freeze-thaw cycle, the autolysis strain was grown, harvested, washed, and resuspended in S30A lysate preparation buffer. The cells were then frozen at −80°C, thawed, vortex mixed, and incubated for 90 minutes at 37°C. The resulting suspension was cleared by centrifugation, the supernatant was stored at −80°C and subse-quently used for transcription-translation (TX-TL) reactions (Methods). Autolysates made by this method were used to set up TX-TL reactions using an established protocol, including the composition of CFE additives (amino acids, nucleotides, energy source, etc) (*27*, *28*). At these conditions, about 19 μM of GFP was produced from a strong lambda phage P_R_ promoter at saturating concentrations of plasmid DNA (Figure 2B-C). Since, except for lysate preparation, the TX-TL reaction conditions used here were essentially as described in ref. (*27*, *28*), we directly compared the performance of the autolysate-based CFE system to the performance of a reference CFE system based on ref. (*28*), which is commercially available (MYtxtl, MYcroarray, Ann Arbor, Michigan). The quantity of GFP produced was similar for the two CFE systems (at saturating DNA concentrations; Figure 2C). Compared to the reference CFE system, GFP expression levels in autolysate-based CFE system were significantly higher at lower template DNA concentrations (≈ 3 nM and less). Although we have not investigated the cause of this phenomenon, it could be due to the potentially higher stability of template DNA in the autolysate owing to the lack of endonuclease I (*endA*) in the parental *E. coli* BL21-Gold (DE3) strain; this endonuclease is known to degrade DNA in plasmid preparations. However, other explanations, such as higher relative transcription efficiency of P_R_ promoter in autolysate-based CFE system, could not be excluded. The maximum rate of GFP production (see Figure 2D (inset) for the definition) was higher in autolysate-based CFE system compared to the reference CFE system (Figure 2D). Comparative analysis of expression of GFP and three other model proteins in the two CFE systems using the same expression cassette demonstrated that expression levels in general can be protein and lysate preparation specific (Figures S2 and S3 of Supporting Information). For example, although both the autolysate-based and the reference CFE systems produced comparable amounts of GFP and chloramphenicol acetyltransferase, the expression levels of Renilla luciferase were higher using the reference CFE system, while *E. coli* β-galactosidase expressed noticeably better in the autolysate-based CFE system.

### 2.4 Expression from linear DNA in autolysates in the presence of GamS protein

One of the major advantages of CFE systems is the speed with which new DNA constructs can be tested. Use of linear DNA, usually produced by standard PCR, further reduces the experimental turn-around time (*18*). Additionally, construct production is much more scalable using linear DNA as it avoids laborious and time-consuming molecular cloning steps, and facilitates production of expression constructs directly by standard assembly PCR (Figure 3A). However, to be used directly in *E. coli* cell lysate-based CFE systems, linear DNA must be protected from degradation by exonucleases, mainly RecBCD, which can be achieved by doping cell extract with lambda phage-derived protein GamS (*19*). In autolysate, GamS protection of linear DNA was quantitative in the conditions tested (Figures 3B and S4 of Supporting Information). Specifically, in the presence of GamS protein the time course of GFP expression from the same molar concentration of a circular plasmid or a linear DNA were nearly identical (Figure 3B). In contrast, in the absence of GamS, GFP production from linear DNA was negligible. Similarly, expression levels of the three other model proteins were indistinguishable using circular DNA, or linear DNA in the presence of GamS (Figure S4). GamS protein could be easily produced in sufficient quantities using a simple protocol (Figures S5 and S6 of Supporting Information; see Methods). Interestingly, in CFE systems based on both lysates prepared by traditional hydrodynamic shearing as well as by egg white lysozyme treatment, GamS protection of linear DNA was noticeably less efficient (*15*, *18*). Although not tested here, GamS production can potentially be further simplified by employing the autolysis strain for GamS overexpression: pAD-GamS plasmid is compatible with pAD-LyseR and therefore can be easily co-transformed in the autolysis strain, which in turn is closely related to our current GamS expression host *E. coli* Tuner (DE3). This would allow replacing the ultrasonication step in the GamS production protocol with a freeze-thaw cycle, reducing the processing time and eliminating the need for ultrasonicator.

**Figure 3:**
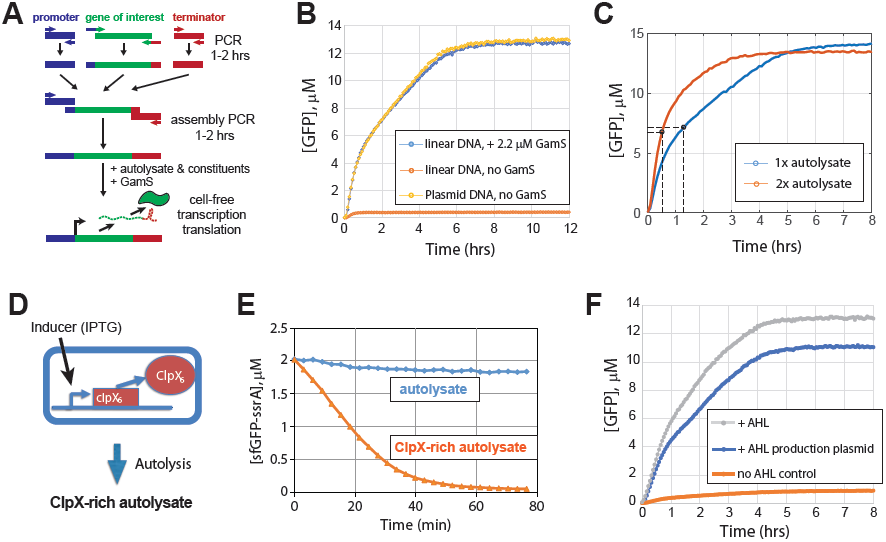
Autolysate validation and applications. (A) Linear DNA gene expression cassettes can be quickly prepared from parts using assembly PCR. A promoter, the gene of interest, and a terminator are amplified by PCR using primers that create overlaps between the parts. The parts are assembled together and amplified in the next round of PCR. The resulting expression cassettes can be directly used in cell-free expression reactions supplemented with GamS protein. (B) Linear DNA is efficiently protected from degradation in autolysate by lambda phage GamS protein. Expression of GFP from a linear PCR product in the presence of GamS (2.2 μM) is indistinguishable from the expression from a circular plasmid at the same concentration (8.1 nM). In comparison, GFP expression from linear DNA in the absence of GamS was negligible. (C) The initial protein production rate can be increased by concentrating the autolysate using standard ultrafiltration. Half-maximum GFP levels are indicated with the black circles. (D, E) Autolysate produced from cells overexpressing an engineered version of ClpX protein has robust ClpXP-protease activity. 2 μM ssrA-tagged GFP was fully degraded with maximum rate of 3.5 μM/hr. (F) Autolysate supports N-acyl homoserine lactone (AHL) production and sensing. All samples contained 7 nM of reporter plasmid with deGFP under AHL inducible *lux* promoter. The samples contained either 130 nM of externally added AHL (+AHL), or 2 nM of N-acyl-homoserine-lactone synthase (*luxI*) producing plasmid (+AHL production plasmid), or neither AHL nor (*luxI*) plasmid (no AHL con-trol).

### 2.5 Concentrated autolysates

We note two modifications to the autolysate protocol that allow the user to control protein synthesis and degradation rates and therefore fine tune gene circuit dynamics, a feature useful for synthetic biology and biotechnology applications. For example, higher production rates may be important for biosensing applications, where the desired sensor output may thus be achieved more quickly. Concentrating the autolysate by ultrafiltration significantly increased protein production rates. Specifically, by concentrating the lysate two-fold, the maximum GFP production rate increased from ≈ 10.2 μM/hr to ≈ 17 μM/hr (Figure S7 of Supporting Information), and half-maximum GFP production was achieved in ≈ 30 minutes compared to ≈ 80 minutes for the parent (1x) autolysate (Figure 3C).

### 2.6 Engineered ClpXP-mediated protein degradation in autolysates

Construction of reversible gene circuits, such as switches or oscillators, requires that their RNA and protein components be degradable. While RNA degradation readily occurs in *E. coli* cell extracts, proteins are typically stable (*29*). In synthetic gene circuits built in live *E. coli* cells, ClpXP protease (reviewed in (*30*)) is often employed to degrade ssrA-tagged proteins and facilitate rapid protein turnover. ClpXP activity is low in unmodified *E. coli* cell lysates possibly because only small quantities of active ClpX protein remain in the cell lysate after preparation (*29*, *31*). An engineered covalently linked ClpX hexamer has been previously purified and added to cell lysates to reconstitute ClpXP activity (*31*, *32*). To prepare lysates with high ClpXP activity “out of the box”, we expressed a covalently linked ClpX hexamer in the autolysis *E. coli* strain at a moderate level (40 μM IPTG induction), which does not measurably affect cell growth (up to 65μM IPTG induction; Figure 3D and Figure S8 of Supporting Information). ClpXP activity was estimated by adding 2 μM sfGFP-ssrA to cell-free reactions supplemented with additional ATP and magnesium (II) (Methods). Autolysate prepared from the ClpX hexamer-overexpressing strain demonstrated ClpXP degradation rate of ≈ 3.5 μM/hr using degradation-tagged superfolder GFP-ssrA as a substrate. TX-TL efficiency of ClpX-rich autolysate was assayed as above for the regular autolysate by quantifying expression of untagged GFP from P_R_ promoter. ClpX-rich autolysate maintained about half of the TX-TL activity in terms of the maximum GFP production rate and the maximum GFP concentration produced (Figure 3E and Figure S9 of Supporting Information). ClpXP-mediated degradation rates can potentially be further increased by concentrating the lysate by ultrafiltration or increasing ClpX hexamer expression in the parent *E. coli* strain.

### 2.7 *Vibrio fischeri* quorum sensing system in autolysates

Many gene circuits used in synthetic biology are based on bacterial quorum sensing systems that utilize N-acyl homoserine lactones (AHLs) for cell-to-cell communication. AHL molecules are produced from acylated acyl-carrier protein (acyl-ACP) and S-adenosyl methionine (SAM) by AHL synthases such as LuxI in *Vibrio fischeri* (*33*). Recently, it has been shown that AHL can be produced within a CFE system by adding luxI encoding plasmid and appropriate precursors (*34*). Based on the idea, we tested if autolysates could produce (and sense) AHL as well. To test it, two plasmids, pLuxI and pLux-deGFP, were constructed. pLuxI plasmid encodes *Vibrio fischeri* N-acyl-homoserine-lactone synthase (LuxI) under a constitutive promoter. pLuxI-deGFP plasmid encodes an AHL sensor: *Vibrio fischeri* LuxR under its native promoter and GFP under *luxI* promoter that is activated by AHL-luxR complex. Our experiments demonstrated efficient activation of *lux* promoter using the AHL producer plasmid pLuxI in autolysate-based CFE system, comparable to its activation with high concentration (130 nM) of externally added AHL (Figure 3F). According to this result, the two AHL precursors, acyl-ACP and SAM, must be present in autolysates. We conclude that, autolysates can be employed in applications that may require quorum based synthetic circuits, such as in pattern formation, collective decision making, and others.

### 2.8 β -Galactosidase-free autolysates

Practical biosensors can benefit greatly from sensitive output that is easy to detect with inexpensive portable equipment or even by naked eye. To this end, we sought to employ a robust and well characterized *E. coli* β-galactosidase enzyme (LacZ), especially since it expressed comparably well in autolysate-based CFE system (Figure S3 of Supporting Information). LacZ is capable of cleaving a number of readily available synthetic substrates resulting in color change, fluorescence, or chemiluminescence, making it easy to adjust the output of LacZ-based biosensors for specific application. For example, the hydrolysis of chlorophenol red-β-D-galactopyranoside (CPRG) results in strong color change from light yellow to red that is easily detectable by naked eye. Similarly, hydrolysis of non-fluorescent fluorescein di-β-D-galactopyranoside (FDG) substrate results in release of strongly fluorescent fluorescein molecule, that can be readily detected. Additionally, chemiluminescent substrates are available for high-sensitivity applications.

Because of high sensitivity of β-galactosidase-based assays, the background levels of this enzyme present in *E. coli* BL21-Gold (DE3) strain result in high β-galactosidase activity in autolysates produced from this strain, essentially precluding its use with LacZ-based readout (Figure 4A). We therefore deleted *lacZ* gene (specifically, the entire lac operon) from the genome of the aforementioned strain, producing *E. coli* BL21-Gold-dLac (DE3). As expected, TX-TL reactions based on autolysates prepared from this strain were devoid of detectable β-galactosidase activity, while retaining their transcription-translation activity (Figure 4A).

**Figure 4:**
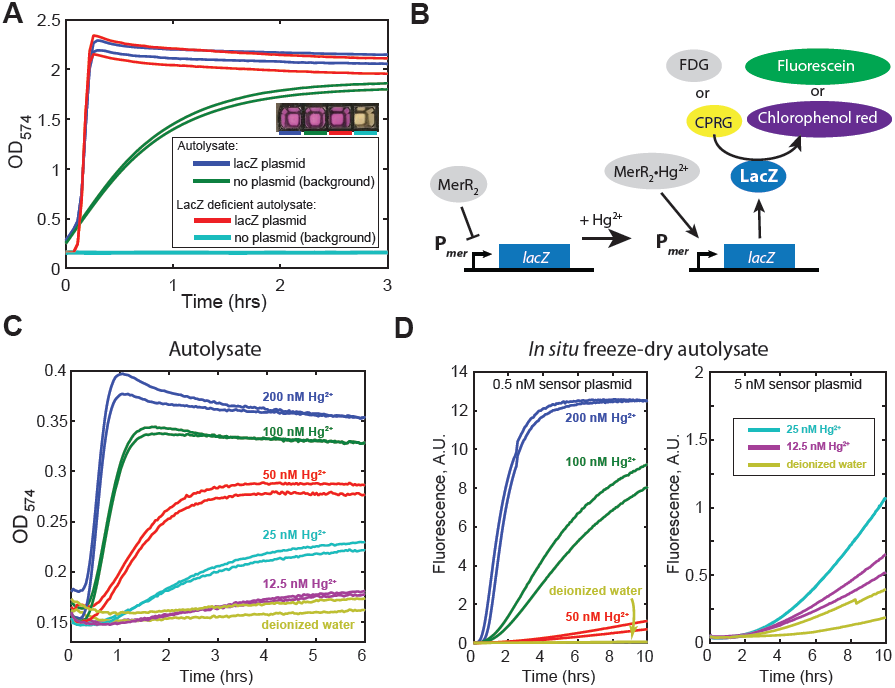
Characterization of β-galactosidase (LacZ) deficient autolysate and construction of LacZ-based mercury (II) sensors. (A) Autolysate produced from an engineered *lacZ-E. coli* strain BL21-Gold-dLac (DE3) does not exhibit detectable β-galactosidase activity. Reactions contained either 2.9 nM pBEST-lacZ plasmid that constitutively expresses LacZ (positive control) or no plasmid. All reactions were supplemented with 300 μM chlorophenol red-β-D-galactopyranoside (CPRG). The results for autolysate produced from BL21-Gold (DE3) strain are shown for comparison. The inset shows a photograph of the reaction wells after overnight incubation at 29°C. (B) Schematic depiction of mercury (II) biosensor-based on *mer* promoter of the mercury resistance operon of Tn21 transposon. In the absence of Hg^2+^P*mer* is repressed by MerR dimer. Upon binding to Hg^2+^MerR dimer changes conformation and activates expression of *E. coli lacZ* gene from *mer* promoter. LacZ enzymatic activity is monitored using chromogenic (CPRG) or fluorogenic (FDC) substrates. (C) The response of autolysate-based Hg^2+^ sensor using chromogenic reporter (300 μM CPRG). (D) The response of Hg^2+^ sensor based on autolysate produced upon rehydration of freeze-dried autolysis strain using fluorogenic reporter (150 μM FDG). Note that the vertical scales on the left and right panels are not directly comparable since the fluorescence was measured with different photomultiplier gains.

### 2.9 Cell-free mercury (II) biosensors

Thus, as a proof of concept we implemented a well-characterized mercury (II) biosensor (*35*– *37*) in CFE systems based on LacZ-free autolysates produced by both freeze-thawing and freeze-drying/rehydration methods (as described above). Briefly, the biosensor was based on mercury resistance operon of transposon Tn21. We constructed a sensor plasmid where P_*merR*_ promoter drives expression of mercury (II) sensor protein, transcriptional regulator MerR, and P_*mer*_ promoter drives MerR regulatable expression of LacZ in the opposite direction (Figure S10 of Supporting Information). In the absence of mercury (II) ions P_*mer*_ promoter is repressed by MerR, however, upon association with mercury (II), MerR activates expression from P_*mer*_ promoter (*38*). Importantly, P_*mer*_ promoter is noticeably leaky in the absence of MerR protein. Thus, MerR protein needs to be present within the TX-TL reaction before the sensor plasmid is added to avoid high background expression of the reporter (LacZ in this case) (*36*). We therefore pre-expressed MerR protein from its native promoter directly within the TX-TL reactions based on freeze-thaw autolysate (45 minutes, 30°C). At this point mercury (II) dilutions as well as the LacZ-based sensor plasmid were added. We detected the output of the reaction using both chromogenic and fluorogenic substrates (Figure 4C and Figure S11 of Supporting Information). With either of the LacZ substrates, the biosensor was sensitive down to at least 12.5 nM mercury (II), eliciting weak but distinguishable response at this concentration, consistently with the previous results (*35*–*37*).

We also implemented the mercury (II) biosensor in CFE system based on LacZ-free autolysates produced upon rehydration of freeze-dried autolysis strain (as described above). In this format, freeze-dried self-lysing *E. coli* BL21-Gold-dLac (DE3) / pAD-LyseR cells supplemented with necessary TX-TL reaction constituents were rehydrated with aqueous solution supplemented with the sensor plasmid (pHg-LacZ), fluorogenic LacZ substrate (FDG), and mercury (II) dilutions. Upon rehydration the cells lyse, starting the TX-TL reaction as described above. To avoid the need to pre-express MerR protein within the TXTL reaction, MerR was expressed in *E. coli* BL21-Gold-dLac (DE3) / pAD-LyseR strain during fermentation using pACYC-MerR plasmid (Figure S12 of Supporting Information; Methods). At the tested conditions, the biosensor in this format had somewhat weaker and less rapid response compared to the same sensor implemented in freeze-thaw autolysate-based CFE systems (compare Figure 4D and Figure S11). However, the sensor was still highly sensitive, eliciting a clear response at 25 nM mercury (II). It is possible that the sensitivity and the response time of the sensor in this format can be further improved, for example by optimizing MerR protein and sensor plasmid concentrations (*36*).

## 3 Discussion

We developed two implementations of the general autolysate production method, using freeze-drying/rehydration and freeze-thawing of cells. The first technique yields dried cells that can produce active lysates in one step by simple rehydration. The second technique produces more traditional cell debris free autolysates. The autolysates prepared this way are compatible with the GamS-based protocol for cell-free expression using linear DNA templates. We demonstrate that protein production rates can be regulated by concentrating the autolysates. At the same time, by preparing autolysates from *E. coli* cells overproducing linked hexameric ClpX protein, autolysates with high intrinsic ClpXP protease activity can be obtained. Without modification, autolysates support N-acyl-homoserine lactone (AHL) biosynthesis and sensing in cell-free expression systems using correspondingly LuxI AHL synthase and LuxR transcriptional regulator from *V. fisheri* quorum sensing system. Finally, autolysates prepared from LacZ deficient cells can be used to implement sensitive background-free LacZ-based output using a variety of substrates, which is illustrated by the implementation of mercury (II) biosensors.

The general approach of producing bacterial lysates for cell-free gene expression using autolysis could facilitate wider adoption of cell-free gene expression systems. Given the modest equipment requirements and simplicity of the preparation protocol it should be accessible to most life sciences laboratories. Specifically, with the possible exception of a high-speed centrifuge, only common equipment is required. No specialized experience or training is required as well, since autolysate preparation protocol relies only on common laboratory procedures. Per reaction cost of autolysate-based cell-free gene expression consists of the cost of producing autolysate as well as the cost of the additional TX-TL reaction components (amino acids, nucleotides, energy regeneration system, etc). The cost of the reaction components (≈ $0.06 per 10 ul TX-TL reaction, not including labor) has been described previously in detail (*27*). This cost heavily depends on the preparation and purchasing scale as well as on the sources of the reagents; for example, we were able to reduce the per reaction cost of amino acids by purchasing them dry instead of pre-dissolved, as in the reference above. If necessary, this cost can likely be further reduced by using less expensive ATP regeneration systems (*28*). The monetary cost of autolysate preparation is comparably low (≈ $0.01 per 10 ul TX-TL reaction), consisting largely of the cost of 2xYT medium. In terms of labor, the protocol takes 8-9 hours from the beginning of fermentation and requires 1 to 2 hours of bench time.

We note a few areas where autolysate production can be potentially improved. First, although we performed the run-off reaction for 90 minutes at 37°C to allow endogenous exonucleases to digest genomic DNA and allow unstable lysate components to precipitate out of solution (Methods; Figure 1), it was previously reported that reducing the duration of this step (or eliminating it completely) significantly improved the performance of lysates based on a *E. coli* B strain, similar to the one used here (*14*). It is therefore possible that run-off duration during autolysate preparation could be reduced, shortening the preparation time and potentially increasing the autolysate activity. Second, in the current protocol, resuspended cell pellets were frozen at −80°C for convenience. It has been recently reported that freezing cells in liquid nitrogen increased the quality of biochemically produced LoFT cell lysate (*15*). Therefore, it is possible that autolysate quality can be increased by liquid nitrogen freezing. Finally, although not investigated here, freeze-drying and rehydration method of producing autolysates can feasibly benefit from additionally expressing phage lambda gene Rz that encodes spanin proteins Rz and Rz1 (Figure S13 of Supporting Information). These proteins aid in disrupting both inner and outer membranes of *E. coli* once the inner membrane is compromized (*23*, *39*), and therefore could facilitate rapid lysate release upon rehydration.

One potential limitation of freeze-dry/rehydration method when applied to biosensor production is the risk that some *E. coli* cells could survive the freeze-drying/rehydration process. In our experiments we found that the typical TX-TL reactions prepared by freeze-drying and rehydration from ≈ 2 mg of cell pellet (≈ 5 *·* 10^9^ colony forming units, CFU) commonly result in 5-50 (occasionally up to 600) CFUs within the reaction (*≈* 10^−9^ *-* 10^−7^ survival rate). This issue will need to be carefully addressed should the sensors be used outside of the laboratory setting. To this end, adjusting lyophilization conditions (*40*), increasing the intracellular lysis protein levels, and adding bactericidal agents that do not interfere with cell-free expression could feasibly be employed, however a careful further investigation of the issue is required.

Although in this paper we focused on production of lysates for cell-free expression, they could also be employed for other emerging synthetic biology and biotechnology applications, such as cell-free proteomics (*4*), metabolic engineering (*1*) or natural product and natural-like product biosynthesis (*8*–*10*). Given the pace of development of cell-free synthetic biology and biotechnology, and the rapidly growing number of researchers working in these fields, we believe that this accessible method of lysate preparation will be of benefit to a large number of researchers.

## 4 Methods

### Cell strains, plasmids, and reagents

All plasmids were constructed using standard PCR and Gibson assembly (*41*) (see Supporting Information for the plasmid DNA sequences). Plasmids pAD-LyseR and pAD-GamS will be available from AddGene: #99244 (pAD-LyseR in *E. coli* BL21-Gold (DE3)), #99245 (pAD-LyseR in *E. coli* BL21-Gold-dLac (DE3)), and #99246 (pAD-GamS in *E. coli* Tuner (DE3)). Unless otherwise specified, the molecular cloning steps were performed in *E. coli* DH5αZ1 (*42*) and *E. coli* Mach1 cells (ThermoFisher Scientific). All plasmids using lambda phage P_R_ promoter (pBEST-variants) were constructed and propagated in *E. coli* KL740 (Yale University Coli Genetic Stock Center, #4382) at 30°C to keep the P_R_ promoter repressed. Where required, antibiotics ampicillin, chloramphenicol, and/or kanamycin were added to cell growth media at 50 μg/ml, 34 μg/ml, and 50 μg/ml respectively. The reference *E. coli* S30 lysate-based transcription-translation system, based on ref. (*28*), was purchased from MYcroarray (Ann Arbor, Michigan; #MYtxtl-70-96-M).

### *E. coli* genome editing

To produce the modified strain *E. coli* BL21-Gold-dLac (DE3), the lac operon in *E. coli* BL21-Gold (DE3) was replaced with spectinomycin cassette from *E. coli* DH5αZ1 (*42*) using pSIM9 helper plasmid according to an established protocol (*43*). Spectinomycin resistance cassette was PCR amplified from E. coli DH5αZ1 using the following primers: 5’-tgcggctaatgtagatcgctgatcggcttgaac-gaattgttagaca, 5’-gaagagagtcaattcagggtggtgatgaaggcacgaacccagt. Homologous arms were PCR amplified from *E. coli* BL21-Gold (DE3) using two primer pairs 5’-accaccctgaattgactctcttc, 5’-atgattttttattgtgcgctcagtatagga and 5’-tggtcttaatatgcgaccactgct, 5’-tcagcgatctacattagccgca. The final construct was produced by assembling the homologous arms and the spectinomycing resistance cassette using assembly PCR followed by amplification using nested PCR with the primers 5’-ATAAACTTGATGGCGGTGCC and 5’-TGAGCTTTACCCGCAAAGTG (see Supporting Information for the complete sequence). The edited strain *E. coli* BL21-Gold-dLac (DE3) was selected for spectinomycin resistance and chloramphenicol sensitivity (i.e. curing of pSIM9 plasmid). The genomic modification was further confirmed by sequencing. The strain will be available through AddGene (#99247).

### Autolysate preparation by freeze-thawing

Unless otherwise specified, all experiments were performed at room temperature. *E. coli* BL21-Gold (DE3) cells (Agilent Technologies) or the engineered *E. coli* BL21-Gold-dLac (DE3) cells (see above) transformed with pAD-LyseR plasmid (Figure S1 of Supporting Information) were used for autolysate production. Transformed cells were grown at 37°C on LB/ampicillin agar plates. Starter cultures were grown overnight at 37°C from a single colony in LB/ampicillin or 2xYT/ampicillin medium. Production cultures were grown in 2xYTPG medium (see Supplemental Experimental Procedures of Supporting Information) supplemented with ampicillin. Typically, 400 ml of 2xYTPG/ampicillin medium was inoculated with 400 μl of starter culture and grown at 37°C on an orbital shaker at 300 rpm in a 1 liter Erlenmeyer flask. Optical density of the culture (OD_600_: 600 nm wavelength, 1 cm pathlength) was measured periodically. To ensure that the measurements were performed within the spectrophotometer linear response range, the sample was diluted 5-fold with the growth medium before measurement. The cells were harvested by centrifugation for 15 minutes at ≈ 1,800 g at room temperature when OD_600_ of the 5-fold culture dilution reached 0.3. This specific harvesting density was chosen based on ref. (*27*) which we did not optimize further. After centrifugation the supernatant was discarded, the pellet was resuspended in ≈ 45 ml of cold (4-10°C) S30A buffer by vortex mixing (see Supplemental Experimental Procedures of Supporting Information), transferred to a preweighed 50 ml centrifuge tube, and centrifugated as above. The wash step can be optionally omitted to reduce the processing time, although we did not estimate the impact of its omission on autolysate quality; the wash step was included for all the preparations used in this paper. The supernatant was then carefully removed by aspiration to allow accurate measurement of the pellet weight. Typically, 1.3 g pellet was obtained from a 400 ml culture. The pellet was resuspended in two volumes (relative to the pellet weight) of cold S30A buffer supplemented with 2 mM dithiothreitol (DTT); for example, 2.6 ml of buffer was added to 1.3 g pellet. The pellet was thoroughly resuspended by vortex mixing and frozen at −80°C. Frozen cell suspensions were thawed in a room temperature water bath, vortex mixed vigorously for 2-3 minutes to potentially facilitate cell lysis, and incubated at 37°C on an orbital shaker at 300 rpm for ≈ 45 minutes. The suspensions were vortex mixed one more time and incubated as above (≈ 45minutes, 37°C, 300 rpm). The sample was cleared by centrifugation for 45-60 minutes at 30,000 g at 4°C. The 30,000 g high-speed centrifugation step could be replaced by a longer centrifugation step at 21,000 g using a common table-top centrifuge, as observed previously (*14*, *15*). However, the higher centrifugation speeds helped to better compact the pellet, increasing the amount of cell debris-free supernatant that could be collected; thus, the autolysates used in this paper were prepared using 30,000 g centrifugation for 45 minutes. The supernatant was removed by pipette and cleared of any remaining cell debris by brief centrifugation (≈ 3-5 minutes) in a table-top centrifuge (we used ≈ 21,000 g – the maximum speed of our centrifuge). Note, the pellet after either of the centrifugation steps can be loose, therefore supernatant must be pipetted out gently to avoid picking up cell debris; we recommend using clear centrifuge tubes in these steps. The lysate should be largely free of cell debris at this point; in our experience, the last centrifugation step can be safely repeated if any debris is still remaining. The lysate was split into small aliquots and frozen at −80°C or used for cell-free expression directly. In our experience, freezing did not affect the lysate performance. For the lysate concentration experiments described in Figure 3D, ultrafiltration was performed using 3,000 NMWL filter (EMD Millipore Amicon Ultra-0.5 Ultracel-3K #UFC500308).

The quality and the quantity of autolysate produced by the freeze-thaw method are sensitive to the ratio of the weight of the cell pellet and the volume of S30A resuspension buffer used to prepare the cell suspension for freezing. More concentrated autolysates typically have higher transcription-translation activity. Thus, it would be desirable to obtain more concentrated autolysates directly by increasing the concentration of cells during the freeze-thaw autolysis step. However, since genomic DNA remains largely intact after autolysis, using high concentrations of cells leads to high viscosity of the solution after lysis. The lysate then becomes difficult to efficiently clear of cell debris by subsequent centrifugation. We found that a cell pellet to buffer ratio of 1:2 (weight per volume) yields the optimal balance between yield and activity of the autolysate. Because of this, it is also critical to accurately measure the pellet weight and the volume of the S30A resuspension buffer to achieve reproducible results. Normally, lower cell pellet to buffer ratios will yield lower amounts of more active lysate and vice versa. Therefore in specific cases it may be beneficial to optimize this ratio within some range, for example 1:1.8–1:2.5.

### Preparation of ClpX-rich autolysate

*E. coli* BL21-Gold (DE3) cells were sequentially transformed with pACYC-FLAG-dN6-His ((*32*), AddGene #22143) and pAD-LyseR plasmids. The cells were grown in 2xYTPG medium supplemented with ampicillin, chloramphenicol, and 40 μM isopropyl β-D-1-thiogalactopyranoside (IPTG). Autolysate was prepared as described above, except that the cell pellet was washed twice with 90 ml of 1x phosphate-buffered saline (PBS) before the final wash with S30A buffer to ensure sufficient removal of chloramphenicol.

### Cell-free expression

The premix solution for cell-free expression was prepared essentially as described previously (*27*, *28*), except for a four-fold increase in concentration of amino acids (final concentration in the cell-free reaction: 4 mM L-leucine, 5 mM – the other 19 amino acids). It was important to supplement the cell-free reactions with additional 2.5-5 mM magnesium glutamate and 0.5-1.5% (w/v) PEG 8000 (final concentrations in cell-free reaction). The specific concentrations of extra magnesium and PEG 8000 should be optimized for new batches of autolysate or premix solutions as non-optimal concentrations of these components can result in poor TX-TL performance. See the detailed protocol in Supplemental Experimental Procedures of Supporting Information. Typically, reactions were set up in a 20 μl volume: 8 μl autolysate + 8.9 μl premix with extra 5 mM magnesium and 1.5% PEG 8000 + 3.1 μl plasmid DNA or other reagents. Unless otherwise noted, the reactions were incubated at 29°C and the progress was monitored by fluorescence measured by microplate reader (Tecan Infinite M200 PRO) using the excitation and emission wavelengths of 485 nm and 520 nm respectively. Purified superfolder GFP was used to calibrate the microplate reader (*44*). Concentrations of deGFP produced within the TX-TL reactions were calculated accounting for its lower relative brightness compared to superfolder GFP (*45*).

For β-galactosidase (LacZ) based readout TX-TL reactions were based on LacZ deficient autolysate produced from *E. coli* BL21-Gold-dLac (DE3) cells. The following β-galactosidase substrates were used: 1) 300 μM chlorophenol red-β-D-galactopyranoside (CPRG) for chromogenic readout or 2) 150 μM fluorescein di-β-D-galactopyranoside (FDG) for fluorescent readout. LacZ-based assays were typically performed at 16 ul scale in microplate reader using 384-well plates with transparent bottom. Chromogenic response (CPRG) was measured by monitoring the optical density of the solution at 574 nm; the fluorescent response (FDG) was measured using the excitation and emission wavelengths of 485 nm and 520 nm respectively.

For SDS-PAGE analysis TX-TL reaction samples were resuspended in an equal volume of loading buffer (100 mM Tris-HCl, pH 6.8, 4% (w/v) sodium dodecyl sulfate (SDS), 0.2% (w/v) bromophenol blue, 20% (v/v) glycerol, 2% β-mercaptoethanol), heated briefly to about 95C, and separated by SDS-PAGE (NuPAGE Novex 10% Bis-Tris Protein Gels, 1.0 mm, 10 well; Invitrogen, NP0301) in MOPS running buffer (Invitrogen, NP0001). The gels were Coomassie stained according to standard procedures. For Western blotting, proteins were transfered to a PVDF membrane (Immobilon-PSQ, Millipore, ISEQ00010) in NuPAGE Transfer Buffer (Invitrogen, NP0006). Staining and development was performed according to standard procedures using the following reagents: anti-6x His tag antibody (primary antibody; Abcam ab9108), anti-rabbit-IgG alkaline phosphatase conjugate (secondary antibody; Sigma, A9919). The staining was performed using 5-bromo-4-chloro-3-indolyl phosphate/nitro blue tetrazolium chromogenic alkaline phosphatase substrate (Sigma B5655).

### Cell-free expression using linear templates

Cell-free gene expression reactions, set up as described above, were supplemented with 2.2 μM GamS protein purified as described below. Linear template was prepared either by assembly PCR (for chloramphenicol acetyltransferase and Renilla luciferase, see below) or by PCR amplification from a previously constructed plasmid (pBEST-LacZ and pBEST-OR2-OR1-Pr-UTR1-deGFP-T500 plasmids) with pBEST-promoter-fw and pBEST-terminator-rev primers (see Supplemental Experimental Procedures of Supporting Information). Linear template DNA was added to cell-free expression reaction last.

### Assembly PCR

All PCR reactions were performed using Q5 High-Fidelity DNA polymerase (New England Biolabs #M0491) according to the manufacturer’s protocol (1x Q5 reaction buffer, 200 μM dNTPs, 0.5 μM each primer, 0.02 U/μl DNA polymerase). 23 PCR cycles were used for all reactions. All reactions were done in 200 μl total volume. The annealing temperature was 60°C. P_R_ promoter and T500 terminator fragments were PCR amplified from pBEST-OR2-OR1-Pr-UTR1-deGFP-T500 plasmid (*28*) using pBEST-promoter-fw/rev and pBEST-terminator-fw/rev primer pairs (see Supplemental Experimental Procedures of Supporting Information). Chloramphenicol acetyltransferase and Renilla luciferase genes were amplified using pBEST-cat-fw/rev and pBEST-Rluc-fw/rev primer pairs from plasmid templates (see Supplemental Experimental Procedures of Supporting Information). Correct PCR product size was confirmed by standard agarose gel electrophoresis. The completed PCR reactions were supplemented with DpnI restriction enzyme (0.4 U/μl, New England Biolabs) and incubated for 1 hr at 37°C to digest the template DNA. The reactions were purified using QIAquick PCR Purification Kit (Qiagen #28104) according to manufacturer’s instructions, eluted in 25 μl of 10 mM Tris-Cl, pH 8.5 (EB buffer), and diluted with EB buffer tenfold. Promoter, terminator, and respective gene fragments were mixed in equimolar amounts and used as a template for the next round of PCR with pBEST-promoter-fw and pBEST-terminator-rev primers. The correct size of the assembled products was confirmed by standard agarose gel electrophoresis. The reactions were purified using QIAquick PCR Purification Kit according to manufacturer’s instructions.

### Expression and purification of GamS

The previously published procedure (*18*) for expression and purification of His-tagged GamS protein was modified in the following way. The entire GamS-6His open reading frame from P_BAD_ promoter-based GamS expression vector P_araBAD-gamS (*18*) was subcloned into pRSF-1b (EMD Millipore) high copy number T7*lac* promoter-based vector, creating pAD-GamS plasmid (Figure S4 of Supporting Information; see Supporting Information for the complete sequence). The plasmid was transformed into *E. coli* Tuner (DE3) cells (EMD Millipore) for expression. 400 ml of LB/kanamycin was inoculated with a starter culture and grown to OD_600_ ≈ 0.4 at 37°C on an orbital shaker. The cell culture was cooled down to room temperature in a water bath, induced with 300 μM IPTG, and grown in an orbital shaker at 150 rpm at room temperature (≈ 23-24°C) for 21 hours. The cells were harvested by centrifugation at 3,400 g for 15 minutes, and stored at −80°C. Frozen pellets were resuspended in 20 ml of GamS lysis buffer (see Supplemental Experimental Procedures of Supporting Information) and resuspended by vortex mixing. The cells were lysed by ultrasonication (Branson Digital Sonifier, 70% power, 5 seconds power on, 30 seconds power off, 20 cycles total, performed on ice). The solution was cleared by centrifugation (25 minutes, 30,000 g, 4°C). Supernatant was applied to Ni-NTA resin using gravity flow (0.5 ml Ni-NTA agarose, Qiagen #30210) and the flow-through was reapplied to the resin one more time. The resin was washed with 20 ml GamS wash buffer and eluted with GamS elution buffer (see Supplemental Experimental Procedures of Supporting Information). The fractions were analyzed by standard Coomassie Blue stained SDS-PAGE (Figure S5 of Supporting Information). Elution fractions containing GamS protein were combined (approximately 3ml total volume) and dialyzed against 500 ml of GamS dialysis buffer (see Supplemental Experimental Procedures of Supporting Information) at 4°C overnight using 3,500 Da molecular weight cutoff dialysis membrane (ThermoFisher Scientific Slide-A-Lyzer #66330). The sample solution contained small amount of precipitate next day and therefore was cleared by centrifugation (16,100 g, 15 minutes, 4°C). Because of sufficiently high purity of GamS in the supernatant and to streamline the purification protocol, the size exclusion chromatography step of the published GamS purification protocol (*18*) was not performed. The supernatant was directly concentrated by ultrafiltration at room temperature using 3,000 NMWL concentrator (EMD Millipore Amicon Ultra-15 Ultracel-3K #UFC900308). The final concentration of GamS was 2.8 mg/ml in the total volume of ≈ 0.6 ml. The concentration was estimated by light absorbance at 280 nm using the extinction coefficient of 11,460 M^−1^ cm^−1^. GamS solution was aliquoted and stored at −80°C.

### ClpX-rich autolysate activity assay

The reactions were set up as described above for cell-free expression, with following modifications. Since ClpXP requires considerable amount of ATP for its activity, it was important to supplemented the reactions with 3 mM additional ATP (pH 7.2) and 4.5 mM magnesium glutamate (partly to compensate for the magnesium ions chelated by the additional ATP (*46*)) to allow degradation to go to completion. 8 nM pBEST-OR2-OR1-Pr-UTR1-deGFP-T500 plasmid was used for production assays. 2 μM ssrA degradation-tagged superfolder GFP was used in the degradation assays.

### AHL production in autolysate

To validate AHL production in autolysates, two plasmids, pLuxI and pLux-deGFP, were constructed (see Supporting Information for plasmid sequences). pLuxI plasmid encodes *Vibrio fischeri* N-acyl-homoserine-lactone synthase (LuxI) under constitutive promoter. The plasmid backbone is based on pNCS-mTFP1 (Allele Biotechnology, San Diego, CA). pLuxI-deGFP plasmid encodes an AHL sensor: *Vibrio fischeri* LuxR under its native promoter and deGFP under *luxI* promoter that is activated by AHL-luxR complex. Transcription-translation reactions were set up as described above and monitored at 29°C by GFP fluorescence.

### Preparation of *in situ* autolysates by freeze-drying followed by rehydration

Unless otherwise specified, all experiments were performed at room temperature. *E. coli* BL21-Gold (DE3) or LacZ deficient *E. coli* BL21-Gold-dLac (DE3) cells transformed with pAD-LyseR plasmid (Figure S1 of Supporting Information) were used. The cells were grown and harvested in S30A buffer with 2 mM DTT as described above for autolysate preparation by freeze-thawing. As above, the pellet was resuspended in two volumes (relative to the pellet weight) of cold S30A buffer supplemented with 2 mM DTT; for example, 2.6 ml of buffer was added to 1.3 g pellet. The pellet was thoroughly resuspended by vortex mixing. Then, 8 ul of the cell suspension was mixed with 8.9 ul of TX-TL reaction premix with extra magnesium and PEG8000 (the same as for the freeze-thaw autolysate reactions above, see Supplemental Experimental Procedures of Supporting Information) and 3.1 ul of either deionized water or plasmid solution in EB buffer. 16 ul of this mixture was transferred into a well of a 384-well microplate and freeze-dried using a commercial freeze-drier (Harvest Right, North Salt Lake, Utah). Freeze-dried samples were typically used within a week of room temperature storage. The reactions were subsequently rehydrated with 14 ul of deionized water. Reactions were then incubated at 29°C and the progress was monitored by GFP fluorescence measured by microplate reader (Tecan Infinite M200 PRO). deGFP concentration was estimated as described above for freeze-thaw autolysate. For β-galactosidase (LacZ) based readout 150 μM fluorescein di-β-D-galactopyranoside (FDG) was added to the reactions. LacZ-based assays were performed at 14 ul scale in microplate reader using 384-well plates with transparent bottom. The cells and premix were directly freeze-dried in the wells beforehand. Fluorescent response (FDG) was measured using the excitation and emission wavelengths of 485 nm and 520 nm respectively.

### Mercury (II) biosensors

Freeze-thaw autolysate-based transcription-translation reactions were set up using LacZ deficient autolysate as described above. Either 300 μM CPRG or 150 μM FDG were added to the reactions for chromogenic or fluorogenic readout respectively. Since even in the absence of mercury (II) ions *mer* promoter is leaky when MerR transcriptional regulator is not present, to significantly reduce the background responce of the sensor in the absence of mercury (II), MerR protein was preexpressed from 2 nM pHg-deGFP plasmid within TX-TL reactions (45 minutes at 30°C). Mercury (II) dilutions and required concentrations of pHg-lacZ sensor plasmid were then added to the TX-TL reactions. The progress of the reactions was monitored either by optical density or fluorescence as described above.

To produce mercury (II) sensor based on autolysates produced from freeze-dried cells, *E. coli* BL21-Gold-dLac (DE3) strain was transformed with pAD-LyseR and pACYC-MerR plasmids. pACYC-MerR plasmid expresses MerR transcriptional regulator from T7*lac* promoter. To induce MerR expression, 20 μM IPTG was added to the medium during cell growth. The cells were grown, harvested, and freeze-dried as described above. To initiate cell autolysis followed by TX-TL reaction the cells were rehydrated with aqueous solution containing the desired mercury (II) dilutions, 0.5 or 5 nM pHg-lacZ sensor plasmid, and 150 μM FDG fluorogenic LacZ substrate. The reaction progress was measured using the excitation and emission wavelengths of 485 nm and 520 nm respectively.

## 5 Associated content

### Supporting information

Supporting-information.pdf: Figures S1-S13;

Supplemental-experimental-procedures.pdf: Supplemental Experimental Procedures plasmid-sequences.zip: plasmid sequences;

## Acknowledgement

The authors thank Omar M. Din, Robert Cooper, Henrike Niederholtmeyer, and Ti Wu for discussions and comments on the manuscript; Simpson Joseph and Xinying Shi for discussions and sharing purified GFP variants; Leo Baumgart and Ryan Johnson for providing the initial mercury biosensor plasmid design; Zachary Sun and Richard Murray for providing P_araBAD-gamS plas-mid. This work was supported by the ARO MURI and the National Institutes of Health grants.

## Graphical TOC Entry

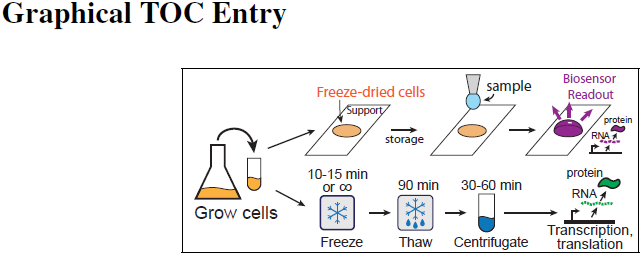

## References

1. Dudley, Q. M., Karim, A. S., and Jewett, M. C. (2015) Cell-free metabolic engineering: biomanufacturing beyond the cell. Biotechnology journal 10, 69–82.

2. Smith, M. T., Wilding, K. M., Hunt, J. M., Bennett, A. M., and Bundy, B. C. (2014) The emerging age of cell-free synthetic biology. FEBS letters 588, 2755–2761.

3. Casteleijn, M. G., Urtti, A., and Sarkhel, S. (2013) Expression without boundaries: cell-free protein synthesis in pharmaceutical research. International journal of pharmaceutics 440, 39– 47.

4. He, M. (2008) Cell-free protein synthesis: applications in proteomics and biotechnology. New biotechnology 25, 126–132.

5. Pardee, K., Green, A. A., Ferrante, T., Cameron, D. E., DaleyKeyser, A., Yin, P., and Collins, J. J. (2014) Paper-based synthetic gene networks. Cell 159, 940–954.

6. Pardee, K. et al. (2016) Rapid, low-cost detection of Zika virus using programmable biomolecular components. Cell 165, 1255–1266.

7. Pardee, K., Slomovic, S., Nguyen, P. Q., Lee, J. W., Donghia, N., Burrill, D., Ferrante, T., McSorley, F. R., Furuta, Y., Vernet, A., Lewandowski, M., Boddy, C. N., Joshi, N. S., and Collins, J. J. (2016) Portable, On-Demand Biomolecular Manufacturing. Cell 167, 248–259.

8. Goering, A. W., Li, J., McClure, R. A., Thomson, R. J., Jewett, M. C., and Kelleher, N. L. (2016) In vitro reconstruction of nonribosomal peptide biosynthesis directly from DNA using cell-free protein synthesis. ACS synthetic biology 6, 39–44.

9. Bashiruddin, N. K., and Suga, H. (2015) Construction and screening of vast libraries of natural product-like macrocyclic peptides using in vitro display technologies. Current opinion in chemical biology 24, 131–138.

10. Maini, R., Umemoto, S., and Suga, H. (2016) Ribosome-mediated synthesis of natural product-like peptides via cell-free translation. Current opinion in chemical biology 34, 44– 52.

11. Takahashi, M. K., Hayes, C. A., Chappell, J., Sun, Z. Z., Murray, R. M., Noireaux, V., and Lucks, J. B. (2015) Characterizing and prototyping genetic networks with cell-free transcription–translation reactions. Methods 86, 60–72.

12. Siegal-Gaskins, D., Tuza, Z. A., Kim, J., Noireaux, V., and Murray, R. M. (2014) Gene circuit performance characterization and resource usage in a cell-free “breadboard”. ACS synthetic biology 3, 416–425.

13. Caschera, F., and Noireaux, V. (2014) Integration of biological parts toward the synthesis of a minimal cell. Current opinion in chemical biology 22, 85–91.

14. Kwon, Y.-C., and Jewett, M. C. (2015) High-throughput preparation methods of crude extract for robust cell-free protein synthesis. Scientific reports 5.

15. Fujiwara, K., and Doi, N. (2016) Biochemical preparation of cell extract for cell-free protein synthesis without physical disruption. PloS one 11, e0154614.

16. Shrestha, P., MichaeleHolland, T., and Charles Bundy, B. (2012) Streamlined extract preparation for Escherichia coli-based cell-free protein synthesis by sonication or bead vortex mixing. Biotechniques 53, 163.

17. Fujiwara, K., and Shin-ichiro, M. N. (2013) Condensation of an additive-free cell extract to mimic the conditions of live cells. PloS one 8, e54155.

18. Sun, Z. Z., Yeung, E., Hayes, C. A., Noireaux, V., and Murray, R. M. (2013) Linear DNA for rapid prototyping of synthetic biological circuits in an Escherichia coli based TX-TL cell-free system. ACS synthetic biology 3, 387–397.

19. Sitaraman, K., Esposito, D., Klarmann, G., Le Grice, S. F., Hartley, J. L., and Chatterjee, D. K. (2004) A novel cell-free protein synthesis system. Journal of biotechnology 110, 257–263.

20. Salehi, A. S., Smith, M. T., Bennett, A. M., Williams, J. B., Pitt, W. G., and Bundy, B. C. (2016) Cell-free protein synthesis of a cytotoxic cancer therapeutic: Onconase production and a just-add-water cell-free system. Biotechnology journal 11, 274–281.

21. Smith, M., Berkheimer, S., Werner, C., and Bundy, B. (2014) Lyophilized Escherichia coli-based cell-free systems for robust, high-density, long-term storage. BioTechniques 56, 186– 193.

22. Young, R. (2013) Phage lysis: do we have the hole story yet? Current opinion in microbiology 16, 790–797.

23. Garrett, J., Fusselman, R., Hise, J., Chiou, L., Smith-Grillo, D., Schulz, J., and Young, R. (1981) Cell lysis by induction of cloned lambda lysis genes. Molecular and General Genetics MGG 182, 326–331.

24. Crabtree, S., and Cronan, J. E. (1984) Facile and gentle method for quantitative lysis of Escherichia coli and Salmonella typhimurium. Journal of bacteriology 158, 354–356.

25. Garrett, J.M., and Young, R. (1982) Lethal action of bacteriophage lambda S gene. Journal of virology 44, 886–892.

26. Kim, R., and Choi, C. (2000) Expression-independent consumption of substrates in cell-free expression system from Escherichia coli. Journal of biotechnology 84, 27–32.

27. Sun, Z. Z., Hayes, C. A., Shin, J., Caschera, F., Murray, R. M., and Noireaux, V. (2013) Protocols for implementing an Escherichia coli based TX-TL cell-free expression system for synthetic biology. JoVE (Journal of Visualized Experiments) e50762–e50762.

28. Shin, J., and Noireaux, V. (2010) Efficient cell-free expression with the endogenous E. coli RNA polymerase and sigma factor 70. Journal of biological engineering 4, 1.

29. Garamella, J., Marshall, R., Rustad, M., and Noireaux, V. (2016) The all E. coli TX-TL Tool-box 2.0: a platform for cell-free synthetic biology. ACS synthetic biology 5, 344–355.

30. Baker, T. A., and Sauer, R. T. (2012) ClpXP, an ATP-powered unfolding and protein-degradation machine. Biochimica et Biophysica Acta (BBA)-Molecular Cell Research 1823, 15–28.

31. Sun, Z., Kim, J., Singhal, V., and Murray, R. M. (2015) Protein degradation in a TX-TL cell-free expression system using ClpXP protease. bioRxiv http://biorxiv.org/content/early/2015/05/22/019695.

32. Martin, A., Baker, T. A., and Sauer, R. T. (2005) Rebuilt AAA+ motors reveal operating principles for ATP-fuelled machines. Nature 437, 1115–1120.

33. Schaefer, A. L., Hanzelka, B. L., Parsek, M. R., and Greenberg, E. P. (2000) Detection, purification, and structural elucidation of the acylhomoserine lactone inducer of Vibrio fischeri luminescence and other related molecules. Methods in enzymology 305, 288–301.

34. Lentini, R., Martín, N. Y., Forlin, M., Belmonte, L., Fontana, J., Cornella, M., Martini, L., Tamburini, S., Bentley, W. E., and Jousson, O. (2017) Two-way chemical communication between artificial and natural cells. ACS central science 3, 117–123.

35. Ivask, A., Hakkila, K., and Virta, M. (2001) Detection of organomercurials with sensor bacteria. Analytical chemistry 73, 5168–5171.

36. Pellinen, T., Huovinen, T., and Karp, M. (2004) A cell-free biosensor for the detection of.transcriptional inducers using firefly luciferase as a reporter. Analytical biochemistry 330, 52– 57.

37. Virta, M., Lampinen, J., and Karp, M. (1995) A luminescence-based mercury biosensor. Analytical Chemistry 67, 667–669.

38. Condee, C. W., and Summers, A. (1992) A mer-lux transcriptional fusion for real-time examination of in vivo gene expression kinetics and promoter response to altered superhelicity. Journal of bacteriology 174, 8094–8101.

39. Berry, J., Summer, E. J., Struck, D. K., and Young, R. (2008) The final step in the phage infection cycle: the Rz and Rz1 lysis proteins link the inner and outer membranes. Molecular microbiology 70, 341–351.

40. Smith, M. T., Bennett, A. M., Hunt, J. M., and Bundy, B. C. (2015) Creating a completely Şcell free Ťsystem for protein synthesis. Biotechnology progress 31, 1716–1719.

41. Gibson, D. G., Young, L., Chuang, R.-Y., Venter, J. C., Hutchison, C. A., and Smith, H. O. (2009) Enzymatic assembly of DNA molecules up to several hundred kilobases. Nature methods 6, 343–345.

42. Lutz, R., and Bujard, H. (1997) Independent and tight regulation of transcriptional units in Escherichia coli via the LacR/O, the TetR/O and AraC/I1-I2 regulatory elements. Nucleic acids research 25, 1203–1210.

43. Sharan, S. K., Thomason, L. C., Kuznetsov, S. G., and Court, D. L. (2009) Recombineering: a homologous recombination-based method of genetic engineering. Nature protocols 4, 206– 223.

44. Pédelacq, J.-D., Cabantous, S., Tran, T., Terwilliger, T. C., and Waldo, G. S. (2006) Engineering and characterization of a superfolder green fluorescent protein. Nature biotechnology 24, 79–88.

45. Day, R. N., and Davidson, M. W. (2009) The fluorescent protein palette: tools for cellular imaging. Chemical Society Reviews 38, 2887–2921.

46. Wilson, J. E., and Chin, A. (1991) Chelation of divalent cations by ATP, studied by titration calorimetry. Analytical biochemistry 193, 16–19.

